# TIP family aquaporins play role in chloroplast osmoregulation and photosynthesis

**DOI:** 10.1101/2020.09.18.297978

**Authors:** Azeez Beebo, Ahmad Zia, Christopher R. Kinzel, Andrei Herdean, Karim Bouhidel, Helmut Kirchhoff, Benoît Schoefs, Cornelia Spetea

**Affiliations:** Department of Biological and Environmental Sciences, University of Gothenburg, Box 461, 405 30 Gothenburg, Sweden; Institute of Biological Chemistry, Washington State University, Pullman, WA, USA; Université de Bourgogne, UMR1347 Agroécologie, ERL CNRS 6300, BP 86510, F-21065 Dijon Cedex, France; MicroMar, Mer, Molécules, Santé EA2160, LUNAM Université, IUML – FR 3473 CNRS, University of Le Mans, 72085 Le Mans Cedex 9, France

**Keywords:** chloroplast, aquaporin, tonoplast intrinsic protein, osmoregulation, photosynthesis, *Arabidopsis thaliana*

## Abstract

Photosynthetic oxygen evolution by photosystem II requires water supply into the chloroplast to reach the thylakoid lumen. A rapid water flow is also required into the chloroplast for optimal oxygen evolution and to overcome osmotic stress. The mechanisms governing water transport in chloroplasts are largely unexplored. Previous proteomics indicated the presence of three aquaporins from the tonoplast intrinsic protein (TIP) family, TIP1;1, TIP1;2 and TIP2;1, in chloroplast membranes of *Arabidopsis thaliana*. Here we revisited their location and studied their role in chloroplasts. Localization experiments indicated that TIP2;1 resides in the thylakoid, whereas TIP1;2 is present in both thylakoid and envelope membranes. Mutants lacking TIP1;2 and/or TIP2;1 did not display a macroscopic phenotype when grown under standard conditions. The mutant chloroplasts and thylakoids underwent less volume changes than the corresponding wild type preparations upon osmotic treatment and in the light. Significantly reduced rates of photosynthetic electron transport were obtained in the mutant leaves, with implications on the CO_2_ fixation rates. However, electron transport rates did not significantly differ between mutants and wild type when isolated thylakoids were examined. Less acidification of the thylakoid lumen was measured in mutants thylakoids, resulting in a slower induction of delta pH-dependent photoprotective mechanisms. These results identify TIP1;2 and TIP2;1 as chloroplast proteins and highlight their importance for osmoregulation and optimal photosynthesis. A third aquaporin, TIP1;1, is present in the chloroplast envelope, and may play role in photosynthesis under excessive light conditions, as based on the weak photosynthetic phenotype of its mutant.

## INTRODUCTION

Oxygenic photosynthetic organisms including plants, utilize water to produce O_2_ through the activity of the water-oxidizing complex (WOC) associated with the photosystem II (PSII) complex. PSII as well as PSI, cytochrome b_6_f (Cytb_6_f), and the ATP synthase are located in the thylakoid membrane within the chloroplast. Since WOC is bound to the lumenal side of PSII, water must be replenished into the thylakoid lumen at a high flux to sustain its activity. Previously, it has been reported that photosynthetic water oxidation is influenced by changes in chloroplast volume, and that the chloroplast stroma adjusts volume in response to osmotic stress (Robinson 1985; McCain 1995). In addition, it has been shown that the thylakoid lumen also adjusts volume in the light and in response to osmotic stress (Cruz *et al*., 2001; Kirchhoff *et al*., 2011). Thus, water oxidation and volume changes in the chloroplast require water movement across envelope and thylakoid membranes.

Water is transported across cell membranes either by free diffusion or by facilitated diffusion through water channels (aquaporins). Aquaporins belong to the major intrinsic protein (MIP) superfamily, which in the model plant *Arabidopsis thaliana* (*Arabidopsis*) has 35 members (Johanson *et al*., 2001). Among *Arabidopsis* MIPs, there are 13 plasma membrane intrinsic proteins (PIPs) and 10 tonoplast intrinsic proteins (TIPs). MIPs and especially TIPs mediate not only the transport of water but also of other small uncharged solutes such as H_2_O_2_, urea, ammonia, and glycerol, and play role in plant development, growth and tolerance to stress (examples of reviews (Gomes *et al*., 2009; Li *et al*., 2013)).

How water molecules enter the chloroplast and reach the thylakoid lumen is largely unexplored. Theoretical calculations indicated low values for the basal water permeability of plant chloroplast envelope and thylakoid membranes, comparable to those of artificial lipid bilayers ((Beebo *et al*., 2013) and refs. therein). In the same paper, it has been hypothesized that simple non-regulated diffusion may sustain water supply at steady state photosynthetic rates. However, to explain the above-described fast osmoregulatory changes in the chloroplast, an increase in water permeability with the help of aquaporins has been hypothesized. Indeed, the fast water exchange and the low activation energy for water transport reported in the literature argue for potential presence of aquaporins in the chloroplast ((Beebo *et al*., 2013) and refs. therein).

TIP1;1, TIP1;2 and TIP2;1 have previously been localized to the tonoplast in *Arabidopsis* roots and leaves using immunoblotting and fluorescence microscopy (Hofte *et al*., 1992; Daniels *et al*., 1996; Hunter *et al*., 2007; Beebo *et al*., 2009; Gattolin *et al*., 2009). Large-scale analyses of the chloroplast proteome of *Arabidopsis* did not find any chloroplast-specific aquaporins. They found instead the three TIPs, but their presence was regarded as a probable tonoplast contamination (Ferro *et al*., 2003; Kleffmann *et al*., 2004). Later on, TIP2;1 was found by proteomics in thylakoid preparations, whereas TIP1;1 was found in the envelope, and TIP1;2 in both types of preparations (Zybailov *et al*., 2008; Ferro *et al*., 2010). Most recent proteomic reports of subfractionated thylakoids found TIP2;1 in the stroma thylakoids, but did not find TIP1;2 in any thylakoid subfractions (Tomizioli *et al*., 2014).

Altogether, the presence of aquaporins remained unclear, and it was necessary to revisit the occurrence of the three TIP family aquaporins in *Arabidopsis* chloroplast membranes by direct visualization of fluorescent-tagged TIPs and western blotting. To study their importance for the chloroplast, we have investigated whether mutants lacking either of these proteins display altered osmoregulation and/or photosynthetic activity in standard and high light (HL) conditions.

## RESULTS

### Revisiting TIPs subcellular location with emphasis on the chloroplast

To test whether the three TIP family aquaporins (TIP1;1, TIP1;2 and TIP2;1) are true chloroplast proteins, we have performed a careful investigation of their location using fluorescence microscopy and western blotting with specific antibodies. The *GFP* gene was fused immediately downstream of the full-length *TIP* cDNAs and transformed into *Arabidopsis* plants under the control of each respective endogenous *TIP* gene promoter (*TIPG* plants). Using confocal laser-scanning microscopy, the tonoplast was found clearly labelled with green fluorescence in the *TIPG* leaves (Figure S1), confirming the previously reported location for the three TIP isoforms (Beebo *et al*., 2009; Gattolin *et al*., 2010). Chloroplasts were not visibly labelled with GFP, probably because chlorophyll (Chl) red autofluorescence was masking GFP fluorescence at the given magnification in intact leaves (Figure S1).

Western blots with anti-GFP antibodies indicated the presence of GFP fusion products in chloroplasts purified by Percoll-gradient from *TIPG* plants, whereas no crossreaction was observed in chloroplasts from non-transformed Col plants (Figure 1a). Western blots with an anti-V-ATPase antibody verified that the analysed chloroplast preparations were free of vacuolar membranes. These observations determined us to analyse these preparations by fluorescence microscopy. TIPG chloroplasts displayed green fluorescence in a punctate pattern (Figure 1b). The signal was strongest for *TIP2;1-GFP* and *TIP1;2-GFP* plants, and weak for the *TIP1;1-GFP* plants. The even weaker signal in the chloroplasts from non-transformed (Col) plants is most likely autofluorescence from phenolic compounds exhibiting a large emission the 500-600-nm range (Galvez *et al*., 1998; Brillouet *et al*., 2013). The difference in signal intensity of the various GFP-TIPs detected in western blot analysis and microscopy is in agreement with the native expression levels of these proteins in the chloroplasts of transgenic lines, as based on the use of native promoters for each construct (see Experimental Procedures).

**Figure 1.**
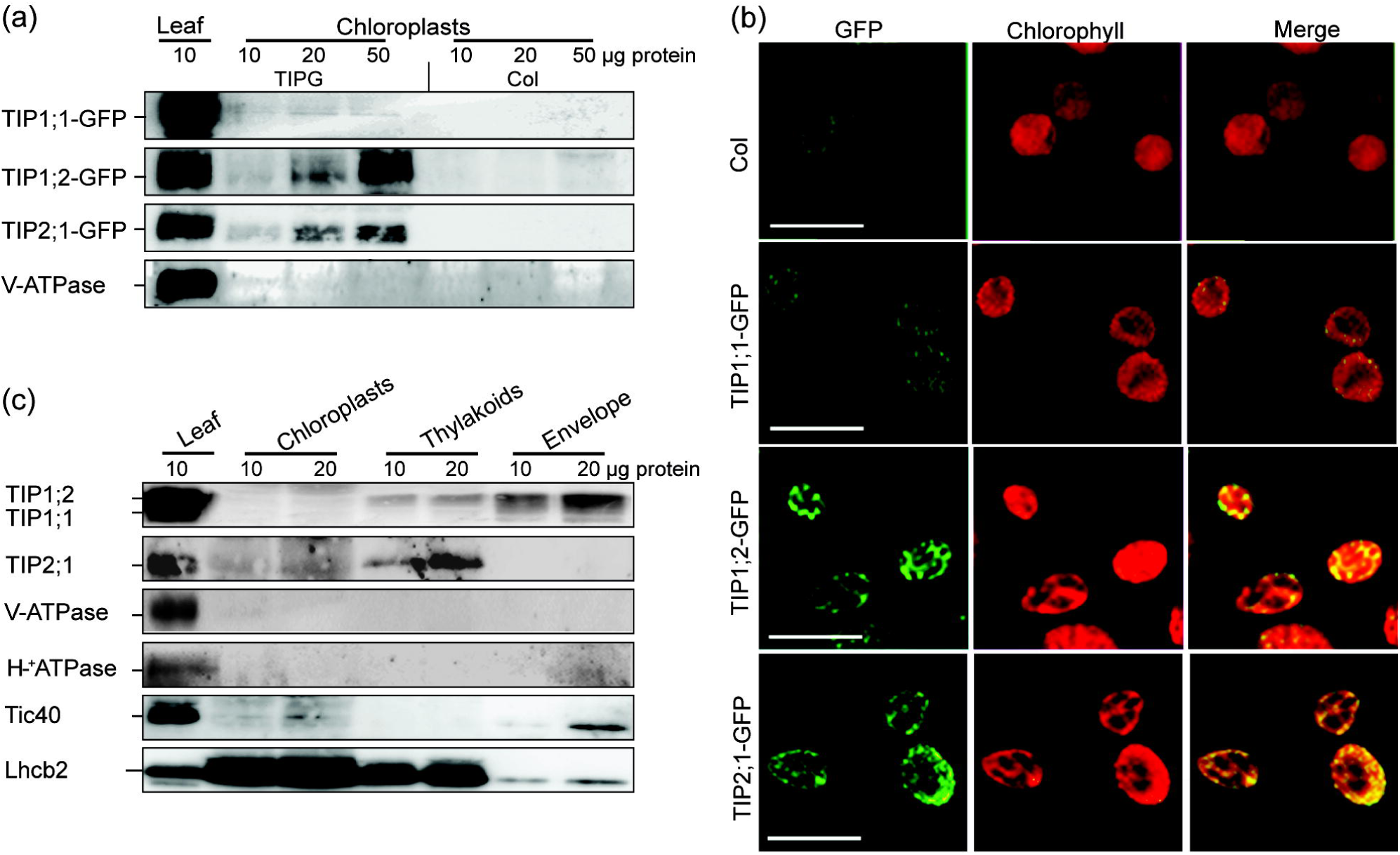
TIPs localization in *Arabidopsis* chloroplasts. (a) Representative Western blots with an anti-GFP antibody of leaf protein extracts and chloroplasts purified from plants expressing TIP-GFP fusion proteins and non-transformed plants (Col). The lack of vacuoles in the analyzed preparations was confirmed using anti-V-ATPase antibodies. (b) Confocal microscopic images of chloroplasts purified from transgenic (TIPG) and Col plants. Scale bar = 10 µm. The fluorescent signal had a punctate distribution in chloroplasts. “*GFP*” represents GFP in the chloroplasts from transformed plants. “*Merge*” shows the overlap of the GFP and the Chl autofluorescence (*CHL*), which was used as a marker for chloroplasts. (c) Representative Western blots with anti-TIP antibodies of leaf extracts, chloroplasts, thylakoids and envelope. The purity of the analyzed preparations was confirmed using anti-V-ATPase, H^+^-ATPase, Tic40 and Lhcb2 antibodies.

Western blot analysis using an antibody against a common peptide for TIP1;1 and TIP1;2 indicated a double polypeptide band in leaf extracts from Col plants (Figure 1c). Purified chloroplasts and also thylakoids displayed the upper polypeptide band. The corresponding band was strongest in the envelope fraction. The lower band of the doublet from leaf extracts was only weakly detected in envelope preparations. The identity of the two bands was resolved by western blotting of leaf extracts from mutants lacking either of the two TIPs, as follows: TIP1;2 as the upper and TIP1;1 as the lower band (Figure 2c). Western blots with an anti-TIP2;1 antibody revealed a single polypeptide band in leaf extracts, also detected in purified chloroplasts and thylakoids. A corresponding band was absent in the envelope. Western blots with antibodies for marker proteins of the tonoplast (V-ATPase), plasma membrane (H^+^-ATPase), inner envelope (Tic40) and the thylakoid membrane (Lhcb2) verified the purity of the tested chloroplast membrane preparations (Figure 1c). Taken together, these results indicate a dual location of the three TIPs, namely in the vacuole and the chloroplast. Within the chloroplast, TIP1;1 and TIP1;2 are present in the envelope, and TIP1;2 and TIP2;1 in thylakoids. TIP1;1 is a well-known tonoplast marker due to its abundance in this membrane (Beebo *et al*., 2009; Gattolin *et al*., 2010), but the fact that the analysed chloroplast preparations were free of tonoplast support a chloroplast location for this TIP as well.

**Figure 2.**
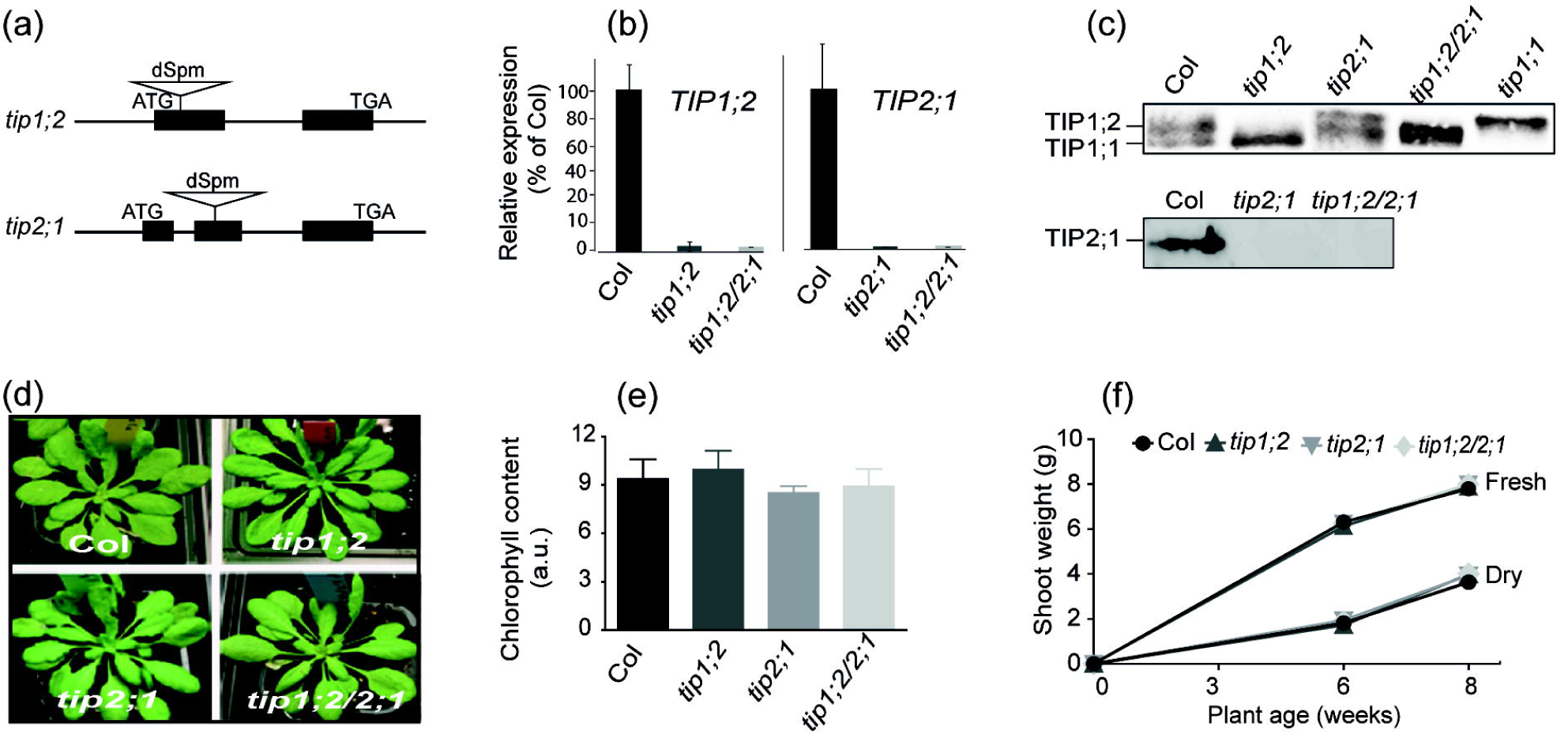
Molecular and growth characteristics of the *tip* mutants. (a) Structure of the Arabidopsis *TIP1;2* and *TIP2;1* genes is shown, indicating the position of the transposon insert (dSpm). (b) Quantitative RT-PCR shows level <3% for *TIP* transcripts in the corresponding genotype. (c) Western blots using a common anti-TIP1;1/TIP1;2 and anti-TIP2;1 antibodies of leaf protein extracts confirmed the lack of the corresponding TIP protein in the respective mutants. (d) Representative photos at 8 weeks under 120 μmol photons m^-2^ s^-1^. (e) Chlorophyll (Chl) content index at an age of 8 weeks. (f) Shoot fresh and dry weight as a function of plant age. The plotted data in (e) and (f) are means ± SD (n = 10-15). No significant differences were observed between the mutants and Col (Student’s t-test, *P* > 0.05).

### Growth phenotype of the *tip* mutants

To investigate the physiological role of the three chloroplast-located TIPs in *Arabidopsis*, single mutants (*tip1;1, tip1;2* and *tip2;1*) and a double mutant (*tip1;2/tip2;1*) carrying either T-DNA or a defective Suppressor-mutator (dSpm) transposable element in the corresponding *TIP* genes (Figure 2a) have been employed. The single *tip1;1* mutant generated in a different background (Ws) than the rest of the lines (Col), was previously described by Beebo *et al*., (2009). Quantitative real-time PCR showed that *TIP1;2* and *TIP2;1* mRNA levels were reduced to < 3% in the respective mutants as compared to Col (Figure 2b). Western blots with TIP1;1/TIP1;2 and TIP2;1 antibodies of leaf extracts indicated the absence of the corresponding protein in the respective *tip* mutant (Figure 2c). The lack of a macroscopic phenotype has previously been reported for the *tip1;1* mutant grown in standard laboratory conditions (Beebo *et al*., 2009). In our study, no differences were observed at an age of 8 weeks in the appearance of the mutants as compared to plants in the respective wild-type background (Figures 2d and S2). Also, no significant differences were obtained in the leaf Chl content, in either the shoot fresh- or dry weight throughout the growth period (Figures 2, e-f and S2, b-c).

### Osmoregulatory volume changes in isolated chloroplasts and thylakoids

We have investigated the effect on osmoregulation of absence of chloroplast TIPs by studying Percoll gradient-purified chloroplasts subjected to osmotic stress for 5 min using a light microscope. When suspended in 0.3 M sucrose (control) buffer, chloroplasts of Col and mutants had an ellipsoid shape and were similar in size (Figure 3). When suspended in water, Col and also *tip2;1* chloroplasts became swollen with irregular shape, whereas those of *tip1;2* and *tip1;2/tip2;1* mutants preserved the ellipsoid shape and size. When incubated in hyper-osmotic solutions containing either ionic (NaCl) or non-ionic (mannitol) osmolyte, all types of chloroplasts decreased in size. Nevertheless, Col and *tip2;1* chloroplasts displayed a much more irregular shape, whereas the *tip1;2* and *tip1;2/tip2;1* mutant chloroplasts were clearly more stable to these osmotic treatments. The lack of osmoregulatory effects in the *tip2;1* chloroplasts is in line with a strict thylakoid location for TIP2;1 (Figure 1c). Thus, the reduced volume changes in the *tip1;2/tip2;1* chloroplasts are most likely due to the absence of TIP1;2 in the envelope. Even though TIP1;1 resides in the envelope, the chloroplasts isolated from the *tip1;1* mutant behaved similarly to those isolated from Ws in all three studied treatments (Figure S3). Taken together, these results strongly indicate that TIP1;2 plays a major role in chloroplast osmoregulation.

**Figure 3.**
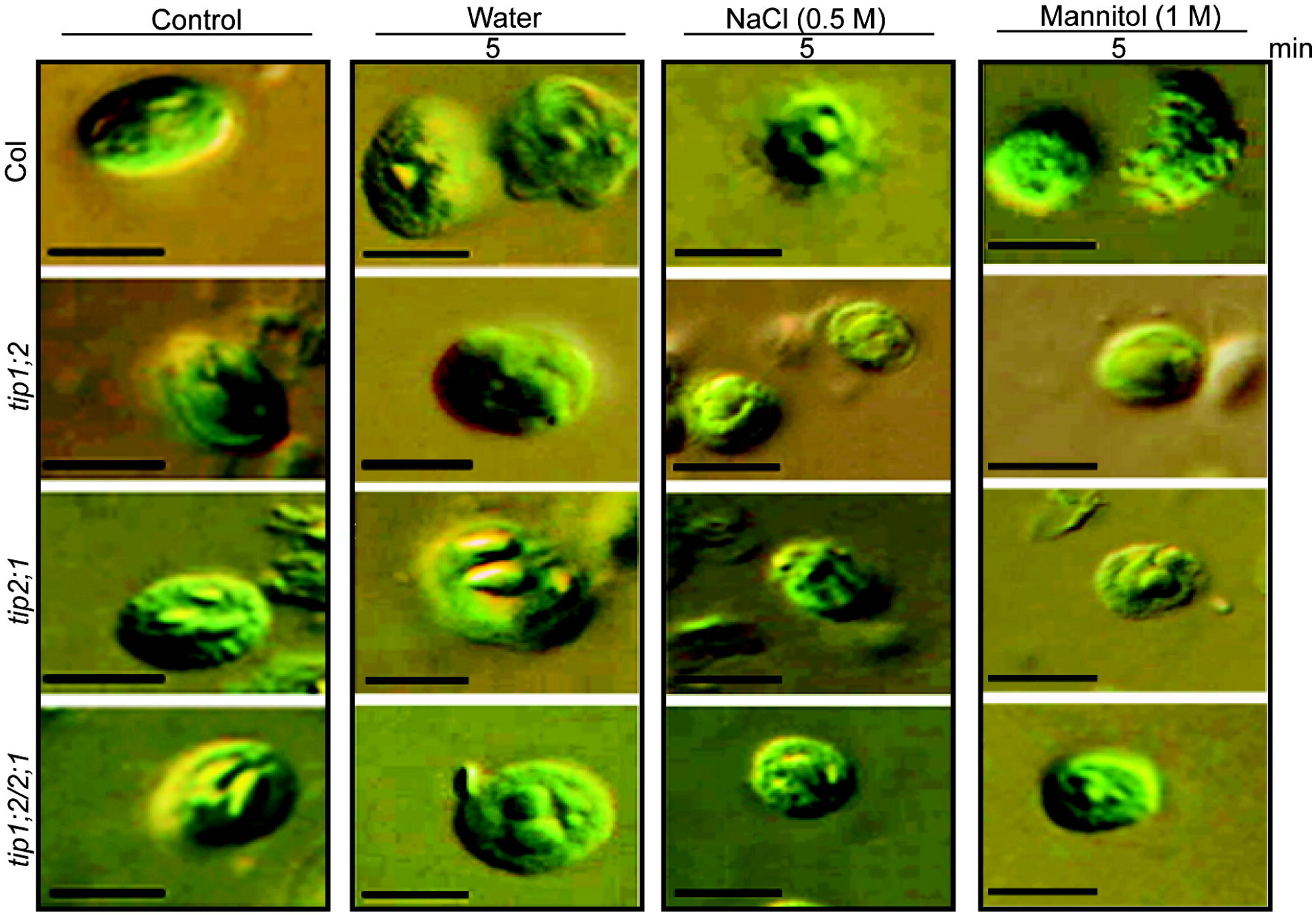
Changes in chloroplast morphology upon hypo- and hyper-osmotic treatments. Light microscopic images were taken on Percoll gradient chloroplasts following a 5 min incubation in 0.3 M sucrose buffer (control), in water, 0.5 M NaCl or 1 M mannitol. Scale bar = 5 μm. The Col and *tip2;1* mutant chloroplasts undergo considerable morphological changes, whereas the *tip1;2* and *tip1;2/tip2;1* mutant chloroplasts appear to be resistant to the osmotic treatments.

Next, we have studied if the lack of TIP1;2 and/or TIP2;1 proteins in Col alters the thylakoid lumen volume changes using 90° light-scattering measurements at 550 nm (ΔA_550,_ Figures 4 and S4). The expansion of the thylakoid lumen in illuminated thylakoids was about 30% smaller (Figure 4 bar plot) in the mutants as compared to Col. Kinetic analysis of light-induced ΔA_550_ revealed that the lower amplitudes in the mutants were caused by about 50% slower formation of the scattering change (Figure 4 *insert*). Next, isolated thylakoids were incubated in sugar-free buffer leading to a swollen state. Starting from this state, addition of sorbitol (hyperosmotic agent) results in water efflux and shrinkage of the thylakoid lumen, as demonstrated previously by electron microscopy (Kirchhoff *et al*., 2011). In Figure 4, the ΔA_550_ values at 300 mM sorbitol were set to zero because this concentration corresponds to the osmolarity of the chloroplast stroma. The sorbitol concentration response curves for all three mutants indicated less osmotic shrinkage of the thylakoid lumen than in Col. In other words, higher concentration of sorbitol is required in the mutants to induce similar osmotic shrinkage as in Col thylakoids.

**Figure 4.**
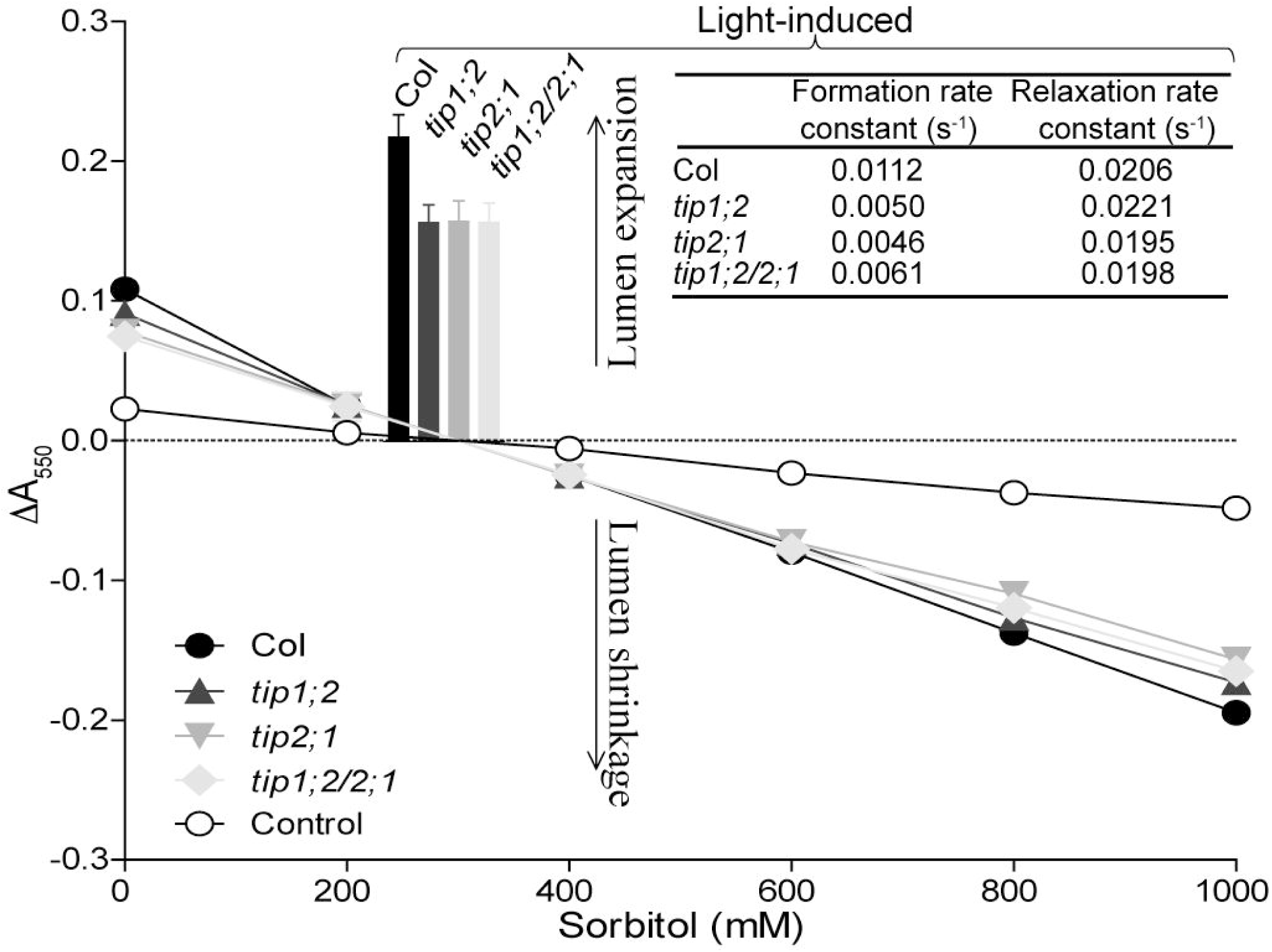
Changes in 90° light scattering in thylakoids during illumination and osmotic treatment. Isolated thylakoids were either illuminated at 1,000 μmol photon m^−2^ s^−1^ in the presence of methyl viologen or incubated in darkness in a buffer containing a series of sorbitol concentrations (for buffer details, see Experimental Procedures). As control, Col thylakoids were incubated in a buffer where the sorbitol was replaced with corresponding volumes of water. Thylakoid lumen volume changes were measured as light scattering change at 550 nm (ΔA_550_). The plotted values are means ± SD (n = 5-7). *Insert*: Formation and relaxation rate constants of the light-induced scattering change.

### Photosynthetic activity in intact leaves and isolated thylakoids

The effect on photosynthetic activity of absence of chloroplast-located TIPs in Arabidopsis was studied in terms of O_2_ evolution, linear electron transport (ETR) and net CO_2_ assimilation (*A*_n_) rates in leaves. Light response curves indicated slight but not significant decrease of O_2_ evolution rate at low-intensity of the photosynthetically active radiation (PAR) in the *tip1;2, tip2;1* and *tip1;2/tip2;1* mutants as compared to Col (Figure 5a *left panel*). At moderate to high PAR intensities (250-2000 µmol m^-2^ s^-1^), the decrease in O_2_ evolution was significantly more pronounced (by up to 30%) in the *tip2;1* and *tip1;2/tip2;1* mutants, indicating the lack of TIP2;1 as the main cause of these effects. Maximal O_2_ evolution was significantly lower in the *tip1;2* and *tip2;1* mutants than in Col, and an additive effect was observed in the *tip1;2/tip2;1* mutant (Figure 5a *right panel*). Similar pattern to the O_2_ evolution was observed for the light response curves of ETR (Figure 5b). The curves for *A*_*n*_ versus leaf intercellular CO_2_ concentration (*C*_*i*_) showed significantly lower assimilation rates in the mutants than in Col at high concentrations (Figure 5c), most likely due to limitation by the reduced ETR rates (Farquhar *et al*., 1980).

**Figure 5.**
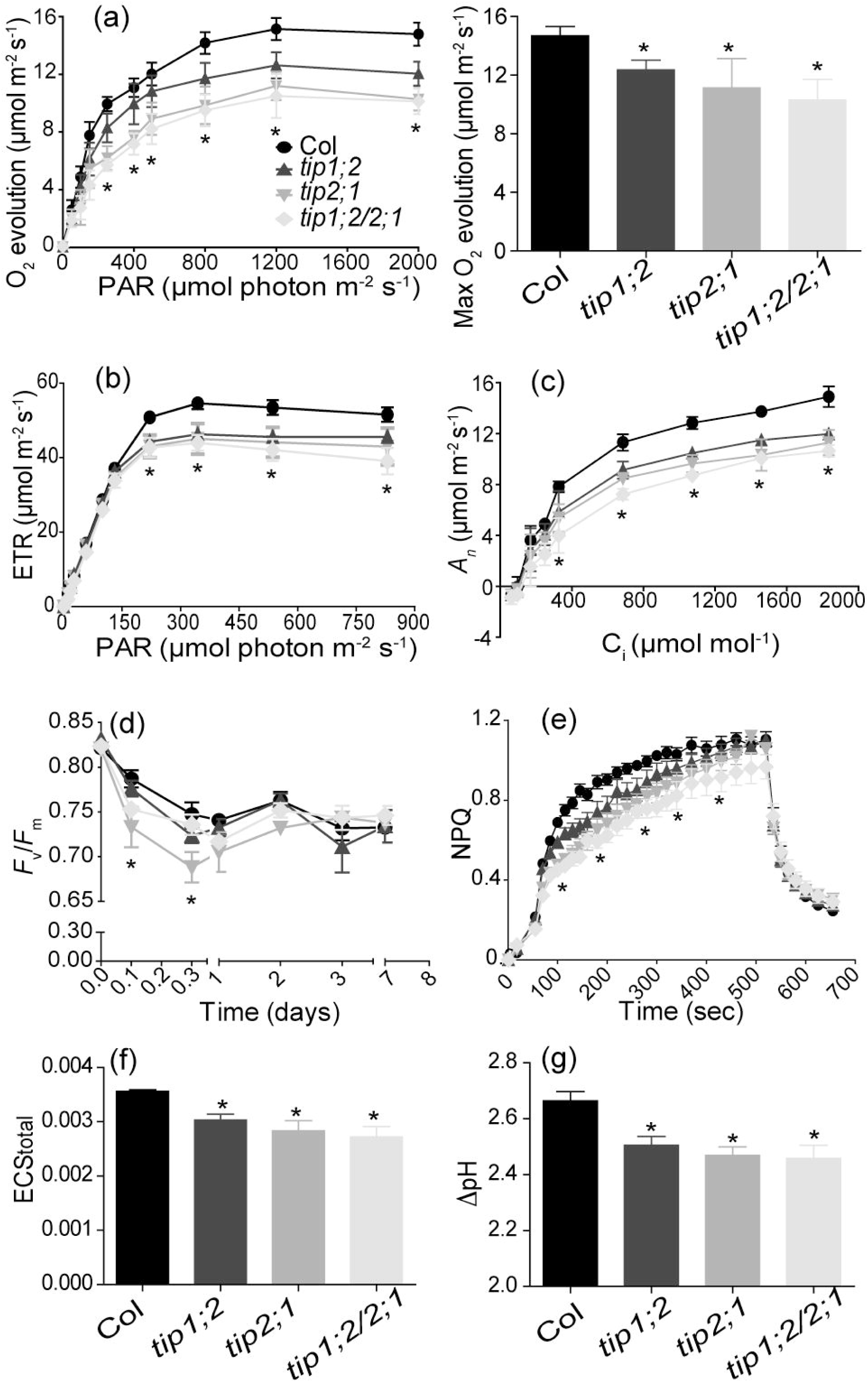
Photosynthetic activity and light sensitivity. (a) *left:* Light response curves of O_2_ evolution rates recorded in leaf discs (n = 3-5). *Right:* Maximal O_2_ evolution from fitted light response curves. (b) Light response curves of electron transfer rates (ETR) recorded on intact plants (n=10). (c) Net CO_2_ assimilation rates (*A*_*n*_) versus intercellular CO_2_ concentrations (*C*_*i*_) recorded in leaf discs using light of 1,800 μmol photon m^-2^ s^-1^ (n = 3-4). (d) *F*_v_/*F*_m_ of intact plants as a function of number of days in high light conditions (800 μmol photon m^-2^ s^-1^) (n = 5). (e) Slow kinetics for induction of non-photochemical quenching (NPQ) formation recorded on intact plants using light of 800 μmol photon m^-2^ s^-1^ (n=3-4). (f) Electrochromic shift (ECS_total_) recorded on intact plants illuminated at 500 μmol photon m^-2^ s^-1^ (n = 5-7). (g) ΔpH recorded as fluorescence of 9-aminoacridine in isolated thylakoids (n = 7-10). The plotted values are means ± SD (a-g). Significantly different values for parameters in the mutants relative to Col are indicated with asterisk (Student’s t-test, *P* < 0.05).

Non-significant effects on various photosynthetic parameters were observed in the *tip1;1* mutant as compared to the Ws background (Figure S5, a-c). Remarkably, steady-state O_2_ evolution in the *tip1;1* mutant became 20% significantly lower than in Ws leaves only at a PAR of about 2,000 μmol m^-2^ s^-1^, indicating that TIP1;1 may play a role in chloroplast photosynthesis only under excessive light conditions. The maximum O_2_ evolution in the *tip1;1* mutant was significantly lower than in Ws leaves (Figure S5a).

Similar abundance of the Lhcb2, D1, AtpB, PsaB, Cytf and RbcL proteins, as determined in western blots with specific antibodies, indicated that the levels of thylakoid photosynthetic complexes and Rubisco were not affected in the mutants as compared to the wild type (Figure S6), and thus cannot explain the observed differences in photosynthetic O_2_ evolution, ETR and CO_2_ fixation rates. Taken together, these results reveal that among the studied mutants, *tip2;1, tip1;2* and *tip1;2/tip2;1* were the most affected in photosynthesis due to the absence of TIP2;1 and TIP1;2 in thylakoids (Figure 2c).

Although mildly, a significantly more pronounced reduction in the maximal photochemical quantum efficiency of PSII (expressed as the Chl fluorescence parameter *F*_*v*_*/F*_*m*_) was observed for the *tip2;1* mutant as compared to Col during 3 h of illumination at a PAR of 600 μmol m^-2^ s^-1^ (HL) (Figure 5d). This indicates an enhanced light sensitivity of the *tip2;1* mutant. However, after 3 days of HL treatment, the *tip2;1* mutant displayed similar *F*_*v*_*/F*_*m*_ as Col. The other mutants were slightly but not significantly more affected than Col throughout the treatment (Figures 5d and S5d). Kinetics of non-photochemical quenching (*NPQ*) formation showed a significantly lower amplitude of the initial rapid phase in the *tip2;1* and *tip1;2/tip2;1* mutants, followed by the *tip1;2* mutant (Figure 5e), which could be attributed to a lower accumulation of H^+^ in the thylakoid lumen as compared to Col (see below). The *tip1;1* mutant displayed only weak but not significant differences from Ws in the *NPQ* kinetics (Figure S5e).

A consequence of reduced PSII and ETR activity in the mutant leaves is that the light-induced H^+^ pumping is reduced, leading to lower proton-motive force (pmf) across the thylakoid membrane. This is indicated by the significantly lower electrochromic shift (ECS_total_) amplitude in leaf measurements (Figure 5f), and supported by ΔpH measurements with 9-aminoacridine (9-AA) in isolated thylakoid membranes (Figure 5g). Taken together, these data indicate that in the light PSII is not fully operational in the *tip1;2, tip2;1* and *tip1;2/tip2;1* mutants, leading to increased photosensitivity and reduced photoprotection.

To study reasons for reduced PSII activity in these mutants, we investigated the photosynthetic reactions in more detail using either intact leaves or isolated thylakoids. Quantification of electron transport components by difference absorption spectroscopy revealed no significant differences in the levels of PSII, Cytb_6_f and PSI complexes between the mutants and Col leaves (Table S1), in accordance with the western blot data (Figure S6). The high-potential chain of electron transport system (between Cytf and P_700_) was probed by analysing relaxation kinetics after a saturating 200-ms light pulse in intact leaves (Kirchhoff *et al*., 2011). At the end of the light pulse, the plastoquinone pool was fully reduced and electron flow into the oxidized high-potential chain was monitored by Cytf^+^ and P_700_^+^ dark relaxation kinetics. Both the unchanged halftime of P_700_^+^ relaxation (Table 1) as well as the redox equilibration between Cytf^+^ and P_700_^+^ (Figure S7) indicated that electron flow in the high-potential chain was unaffected in the three mutants. The *F*_v_/*F*_m_ parameter was indistinguishably high between the mutants and Col leaves (Table 1), indicating that in dark-adapted plants PSII is fully operational upon application of a strong saturation pulse. Notably, the Φ_II_ parameter was significantly lower in the mutant leaves (Table 1), consistent with the light response ETR curves (Figure 5b), indicating that in continuous light PSII does not function properly.

**Table 1.** Photosynthetic parameters in Col and *tip* mutants

| Genotype | $P_{700}^{+} t_{1/2}^a$<br>(ms) | $F_v/F_m$ | $\Phi_{II}$ | O<sub>2</sub> evolution | | |
| --- | --- | --- | --- | --- | --- | --- |
|  |  |  |  | PSII | ETR | Cytb<sub>6</sub>f |
| | | | | $\mu\text{mol O}_2 \text{ mg Chl}^{-1} \text{ h}^{-1}$ | | |
| Col | 6.7 ± 0.8 | 0.8 ± 0.0 | 0.2 ± 0.0 | 423 ± 66 | 215 ± 47 | 695 ± 111 |
|  | 100 ± 11% | 100 ± 0% | 100 ± 12% | 100 ± 16% | 100 ± 22% | 100 ± 10% |
| *tip1;2* | 90 ± 8% | 99.7 ± 1.2% | 75 ± 21% * | 105 ± 21% | 106 ± 22% | 103 ± 22% |
| *tip2;1* | 90 ± 6% | 99.5 ± 0.9% | 75 ± 16% * | 106 ± 18% | 106 ± 22% | 105 ± 20% |
| *tip1;2/2;1* | 90 ± 4% | 99.1 ± 0.9% | 79 ± 8% * | 103 ± 14% | 105 ± 19% | 101 ± 18% |

Plants were grown for 6 weeks and dark-adapted overnight before recording the halftime of $P_{700}^{+}$ ( $t_{1/2}$ ) and $F_v/F_m$ in intact leaves. $\Phi_{II}$ was recorded in intact leaves following illumination. Steady-state rate of PSII-mediated O<sub>2</sub> evolution was measured in isolated thylakoids using dimethyl-benzoquinone as artificial electron acceptor in the presence of the H<sup>+</sup> uncoupler nigericin. Linear electron transfer rate (ETR) was measured using methyl viologen and sodium azide under similar conditions. The Cytb<sub>6</sub>f activity was determined using reduced duroquinol as electron donor and methyl viologen as PSI acceptor. Absolute values are given for Col, whereas mutant data are expressed relative to Col. All values are means ± SD (n = 5-10). Significantly different values within the same column are indicated with asterisk (Student's t-test, $P < 0.05$ ).

When using thylakoids isolated from Arabidopsis Col and mutant leaves, no significant differences were observed for the light-saturated PSII O_2_ evolution rate probed with the electron acceptor dimethyl-benzoquinone in the presence of the uncoupler nigericin (PSII, Table 1). This is in disagreement with the data from leaf measurements (Figure 5a). Furthermore, O_2_ evolution in the presence of methyl viologen (MV, electron acceptor from PSI) and of nigericin revealed no significant reduction for the mutants as compared to Col (ETR, Table 1). Cytb_6_f activity was also indistinguishable between Col and mutants (Cytb_6_f, Table 1). Taken together, these data indicate that water availability is not limiting for the full operational of PSII in thylakoids isolated from the *tip* mutants. Therefore, other factors than water deficiency in the thylakoid lumen may explain the reduced PSII activity in the mutant leaves.

To further explore the role of TIP2;1 in photosynthetic regulation, we performed experiments under fluctuating light. In transition from low to high light, the *tip2;1* mutant displayed slower NPQ induction but without changing the PSII quantum yield (Figure S8). In the TIP2;1-GFP transgenic plants, NPQ was induced faster, resulting in less electron transport through PSII than in Col. These data indicate a role for TIP2;1 in fluctuating light conditions where relatively fast changes in ΔpH across the thylakoid membrane are required. Furthermore, ammonia known to regulate water uptake and aquaporins (Wang et al. 2016) appears to enhance the effects seen in the mutants (Figure S9).

## DISCUSSION

The molecular mechanisms governing water transport into and within the chloroplast are largely unexplored. Nevertheless, recent theoretical considerations based on published experimental data suggested the presence of aquaporins in thylakoids (Beebo *et al*., 2013). In this study, we bring experimental proof for the occurrence of three TIP family aquaporins (TIP1;1, TIP1;2 and TIP2;1) in *Arabidopsis* chloroplast membranes, and also bring insights into their role in chloroplasts. We propose that TIP1;2 and TIP2;1 are housekeeping chloroplast aquaporins, which importantly contribute to its water balance and photosynthesis, whereas TIP1;1 may contribute to the efficiency of photosynthesis only in acute light stress.

### TIP1;1, TIP1;2 and TIP2;1 have dual location to the vacuole and the chloroplast

TIPs are located mainly in the tonoplast, and some of them (*e*.*g*., TIP1;1) have been extensively used as tonoplast markers (Beebo *et al*., 2009; Gattolin *et al*., 2010). Since they do not possess targeting sequences, other subcellular locations could not be ruled out. Proteomics of chloroplast preparations gave contradictory results about the three TIP isoforms (Ferro *et al*., 2010; Tomizioli *et al*., 2014). Therefore, individual localization studies were requested. In this work, green fluorescence microscopy revealed that, in addition to the vacuole, *Arabidopsis* TIP1;1, TIP1;2 and TIP2;1 were associated with the chloroplast as punctate structures (Figures 1b and S1). The fact that in the *TIP1;1-GFP* line, the GFP signal was detected primarily in a few punctate structures near the organelle periphery (Figure 1a) is in line with its low expression level in chloroplasts, as detected by western blotting (Figure 1, a and c). Similar punctate fluorescence pattern has been reported for other chloroplast proteins fused to GFP, namely dynamin FZL, GTPase ObgC, glutamine synthase GLN2, and mechanosensitive channel MSCL1 (Taira *et al*., 2004; Gao *et al*., 2006; Nakayama *et al*., 2007; Bang *et al*., 2009). In the case of ObgC, this pattern was explained by a dimerization-type of protein-protein interaction (Bang *et al*., 2009). Using bimolecular fluorescence complementation in yeast, Murozuka *et al*., (2013) have shown strong interactions between TIP1;2 and TIP2;1, and proposed a role in targeting and/or regulation. FZL was found dually located to the envelope and the thylakoid membrane (Gao *et al*., 2006), GLN2 was found in both the chloroplast stroma and mitochondrial matrix (Taira *et al*., 2004), and MSCL1 was found in the cytoplasm and the chloroplast (Nakayama *et al*., 2007). Thus, the punctate pattern indicates location in distinct areas in the membrane and it may require interaction between TIPs and/or to dual targeting to the vacuole and the chloroplast. Other examples of dually located aquaporins are TIP3;1 and TIP3;2, which were found in the plasma membrane and the tonoplast (Gattolin *et al*., 2011). AQP1 was localized to the plasma membrane and also to the chloroplast envelope in *Nicotiana tabacum* (Uehlein *et al*., 2003; Uehlein *et al*., 2008). Interestingly, the substrate specificity was found to be location-dependent since only the plasma membrane AQP1 transported water, whereas the envelope AQP1 facilitated diffusion of CO_2_. Whether the type of substrate is location-dependent in the case of the three TIPs needs further investigations.

### Chloroplast-located TIPs are important for chloroplast osmoregulation and optimal photosynthesis

Although they may co-exist on the same tonoplast, transcriptomics data and fluorescence microscopy of fusion proteins indicated distinct tissue and organ expression pattern for the three TIPs in *Arabidopsis* (Schmid *et al*., 2005; Hunter *et al*., 2007; Gattolin *et al*., 2009). TIP1;1 is the most widespread TIP isoform in *Arabidopsis*, being present in both roots and leaves (Hunter *et al*., 2007; Beebo *et al*., 2009; Gattolin *et al*., 2009). TIP1;2 and TIP2;1 are mostly expressed in leaves and have only a limited expression in roots (Hunter *et al*., 2007; Gattolin *et al*., 2009). *Arabidopsis tip1;1* and *tip1;2* mutants did not show any macroscopic phenotype in previous reports (Schussler *et al*., 2008; Beebo *et al*., 2009). In this study, in addition to these mutants, *tip2;1* and also *tip1;2/tip2;1* did not display any visual phenotype when grown under standard conditions (Figure 2 and S2). Our and previous observations (Beebo *et al*., 2009) that inactivation of one or another TIP has little or no effect on plant growth under standard conditions indicate redundant functions among this type of aquaporins.

The most beneficial action of aquaporins is to provide massive water flow in response to osmotic stress, leading to rapid changes in cell volume. Indeed, volume changes of the thylakoid lumen and chloroplast stroma have been previously reported upon osmotic shock and also in the light (Robinson 1985; McCain 1995; Cruz *et al*., 2001; Kirchhoff *et al*., 2011). In this study, western blotting using anti-TIP antibodies indicated an envelope location for TIP1;1, a thylakoid location for TIP2;1 and a dual envelope/thylakoid location for TIP1;2 (Figure 1c). Our observations of inhibited osmoregulatory volume changes in the chloroplasts of *tip1;2* and *tip1;2/tip2;1* mutants (Figure 3) indicate a main role for TIP1;2 protein in mediating water movement across the chloroplast envelope. The observations of reduced thylakoid volume changes in the light and also upon osmotic treatment of *tip1;2, tip2;1* and *tip1;2/tip2;1* mutants (Figure 4) point to a role for TIP1;2 and TIP2;1 in water movement across the thylakoid membrane.

Water is a substrate for photosynthetic O_2_ evolution inside the thylakoid lumen. Recent calculations based on published data indicated that free diffusion could sustain water oxidation at steady state photosynthetic rates, and that aquaporins may not be absolutely required (Beebo *et al*., 2013). In this study, we show no difference in photosynthetic O_2_ evolution between Col and the *tip1;2, tip2;1* and *tip1;2/tip2;1* mutants thylakoids (Table 1), consistent with this hypothesis. However, significantly reduced O_2_ evolution rates were obtained in mutant leaves (Figure 4a). There are several possible explanations for this discrepancy. 1) The lack of tonoplast-located TIPs may indirectly affect photosynthesis in intact leaves, but not in isolated thylakoids, since they are devoid of tonoplast (Figure 1c). However, this possibility is not very likely since absence of the most abundant TIP isoform in the tonoplast (TIP1;1) had only marginal effects on photosynthesis in leaves (Figure S5). 2) In isolated thylakoids, water supply into the lumen via free diffusion may sustain PSII photochemistry, making aquaporins not absolutely required, as proposed by Beebo *et al*., (2013). However, aquaporin-dependent processes, such as chloroplast osmoregulation, may limit PSII activity in leaves. Osmoregulatory volume changes were observed in chloroplasts isolated from wild type, *tip1;1* and *tip2;1* but were inhibited in the *tip1;2* and *tip1;2/tip2;1* mutant chloroplasts. Thus, the possibility that defects in vacuole-located TIPs caused the observed effects is excluded. The inhibited osmoregulatory volume changes of chloroplasts isolated from the mutant leaves may reduce the dynamics in thylakoid organization, hence its optimal function in photosynthesis.

The average pre-exchange lifetime of water in the thylakoid lumen of 10 ms indicates that the lumen is refilled 3 × 10^4^ fold faster than the photosynthetic water oxidation when expressed per sec (Beebo *et al*., 2013). Thus, if aquaporins are responsible for this high exchange activity, it is for the optimal function of the thylakoid rather than primarily to sustain water oxidation. This reasoning could explain the mild effects on photosynthetic O_2_ evolution at moderate to high PAR intensities in the *tip1;2, tip2,1* and *tip1;2/tip2;1* mutants (Figure 5). The *tip1;1* mutant was affected in O_2_ evolution only at extremely high PAR, suggesting a role for TIP1;1 under acute light stress in chloroplasts. Finally, Chl fluorescence measurements indicated an earlier HL-induced inactivation and to a larger extent in the *tip2;1* mutant than in Col and the other *tip* mutants (Figure 5), in line with the thylakoid location of TIP2;1. Although significant differences were observed, the effects were relatively mild. One potential explanation could be redundancy of the three studied TIPs and of other aquaporins in the envelope as detected by mass spectrometry (*e*.*g*., PIPs, Ferro *et al*., (2010)).

## CONCLUSIONS

The finding of a chloroplast location in addition to the vacuolar one for the three TIP family aquaporins allowed us to validate the previous proteomics findings. Phenotypic characterization of their mutants allowed us to understand that TIP1;2 and TIP2;1 are necessary for chloroplast osmoregulation and optimal photosynthesis, whereas TIP1;1 may act in extreme light stress conditions. The two roles may be interrelated by a mechanism that requires further investigations. Other questions that should be the focus of future studies are the signals and pathways of TIPs trafficking to the chloroplast. The mechanisms of TIP trafficking in the cytosol are poorly understood, but it is thought that they are delivered to the vacuole by vesicles from the endoplasmic reticulum via Golgi-dependent and independent pathways (Rivera-Serrano *et al*., 2012). Few examples of chloroplast proteins have been reported to use Golgi vesicles (Kitajima *et al*., 2009) and putative components of a vesicular transport system to and within the chloroplast have been predicted (Khan *et al*., 2013). Additional question to be resolved is whether each of the chloroplast TIP isoforms has a specific role or they just provide sufficient redundancy to ensure that cellular water homeostasis remains under control for optimal chloroplast function.

## EXPERIMENTAL PROCEDURES

### Plant material and growth conditions

*Arabidopsis* plants (ecotypes Col and Ws), the *TIPG* transgenic and *tip* mutants were grown using a 16 h dark/8 h light cycle (120 μmol photons m^-2^ s^-1^) at 20°C and 60 % relative humidity) in a growth chamber (CLF PlantMaster, Plant Climatics GmbH, Wertingen, Germany) for 8 weeks, if not otherwise indicated. The *Arabidopsis TIP1;1-GFP* and the *tip1;1* mutant have been previously described (Beebo *et al*., 2009). The construction of the *TIP1*;*2*-*GFP* and *TIP2;1-GFP* fusions and *Arabidopsis* transformation, the *tip1;2, tip2;1* and *tip1;2*/*tip2;1* mutants are described in Methods S1, Tables S2 and S3. For localization studies, chloroplasts were isolated and purified on a Percoll gradient according to (Thuswaldner *et al*., 2007). Following lysis of purified chloroplasts, thylakoids and envelope were purified on a sucrose gradient as described (Thuswaldner *et al*., 2007). Protein concentration was determined using Bio Rad DC™ protein Assay (Bio Rad). For activity studies, chloroplasts and thylakoids were prepared by a fast method at 4°C. In brief, dark-adapted rosette leaves were harvested and homogenized in grinding medium (400 mM sorbitol, 20 mM Tricine, 10 mM EDTA, 10 mM NaHCO_3_ and 0.15% BSA at pH 8.4). The homogenate was then filtered, centrifuged at 2,000 x *g* for 2 min, and the resulting pellet was resuspended in washing medium (400 mM sorbitol, 20 mM HEPES, 2.5 mM EDTA, 5 mM MgCl_2_, 10 mM NaHCO_3_ and 0.15% BSA at pH 7.6). Final pellet was obtained by centrifugation at 2,000 x *g* for 1 min, and resuspended in the washing medium. Chl concentration was measured in 80% (w/v) acetone (Porra 2002). Leaf Chl content was measured using a portable device (CCM-200 plus, Optic-science Inc, NL).

### Microscopy

For localization studies, leaf sections of *TIPG* and Col leaves and Percoll-gradient isolated chloroplasts were mounted in water under a coverslip. Fluorescence detection was conducted with a LSM 700 Axio Observer.Z1 confocal microscope (Carl Zeiss, Germany). Excitation wavelengths and emission filters were 488 nm/band-pass 506–538 nm for GFP and 488 nm/bandpass 664–696 nm for Chl. Images were processed using Photoshop CS5 software (Adobe Systems, San Jose, CA). For osmotic changes, purified chloroplasts were resuspended in control buffer (50 mM Tricine pH 7.5 containing 5 mM MgCl_2_ and 330 mM sucrose), in water, 0.5 M NaCl or 1 M mannitol and incubated for 5 min on ice before examination, as described (Zhang *et al*., 2012). Photos of the chloroplasts were taken with a light microscope in phase contrast mode (Axioplan2 Imaging, Zeiss, Germany).

### Light scattering

Changes in 90° light scattering were measured with a HoribaYvon Fluoromax 4 spectrofluorometer (λ_ex_ = λ_em_ = 550 nm; bandwidth: 2 nm) on isolated thylakoid membranes prepared from dark-adapted plants (Kirchhoff *et al*., 2011). Membranes (20 μg Chl ml^-1^) were suspended in 20 mM Tricine buffer (pH 8.2) containing 150 mM KCl, 7 mM MgCl_2_ and incubated for 10 min to obtain swollen lumen and subsequently incubated for 5 min with sorbitol at various concentrations to induce shrinkage before recording the spectra. These incubation times were initially measured and selected to ensure stable absorbance signal by complete mixing of thylakoid suspension and osmoticum for each measurement. Light scattering changes in the presence of 50 μM MV were induced by applying red light (1,000 μmol m^−2^ s^−1^, Leica KL 1500 LED). Scattering changes were measured first in dark for 100 s, then the thylakoids were light treated for 400 s followed by dark measurement for 500 s. All light-induced scattering changes were measured with continuous stirring while osmoticum induced changes were measured with stirring switched off 2 min prior to each data recording, to avoid signal noise.

### Measurement of photosynthetic activity

Response curves of O_2_ evolution rate versus irradiance were recorded in leaf discs using Clark-type electrode (Hansatech, Pentney, King’s Lynn, U.K.) at 20°C. The curves were fitted according to Nguyen-Deroche Tle *et al*., (2012) using CurveExpert software, to calculate the maximum photosynthetic O_2_ evolution.

Chl *a* fluorescence was measured using a Hansatech FMS1 fluorimeter on intact plants at the end of the 16-h dark phase or following illumination for 6 min with actinic light of 675-725 µmol photon m^-2^ s^-1^. *F*_*v*_*/F*_*m*_ and Φ_II_ were deduced as described (Baker 2008). Light response curves of ETR were determined based on Chl *a* fluorescence measured on intact plants using Dual-PAM-100 (Walz, Effeltrich, Germany) as described (Lundin *et al*., 2007).

*A*_*n*_*/C*_*i*_ response curves were recorded using a leaf gas-exchange instrument (LI-6400XT, LiCOR, Lincoln, Nebraska, USA) at saturating PAR of 1,800 µmol m^-2^ s^-1^ and 25°C, with stepwise increase of external CO_2_ concentration (with 3-5 min of adaptation). For determination of *NPQ*, slow kinetics of Chl fluorescence induction were recorded in intact plants, exposed to PAR of 800 µmol photon m^-2^ s^-1^, followed by darkness.

Steady-state rates of O_2_ evolution were also measured in isolated thylakoids using Clark-type electrode at 20°C. The thylakoids (20 μg Chl ml^-1^) in 25 mM HEPES buffer (pH 7.5), 300 mM sorbitol, 40 mM KCl and 7 mM MgCl_2_ were illuminated with saturating red light (3,000 μmol photon m^-2^ s^-1^) in the presence of 1 μM nigericin. PSII activity was assayed by using 1.5 mM dimethyl-benozoquinone as electron acceptor. Linear ETR was measured with 100 μM MV as the electron acceptor for PSI. MV catalyzes the reduction of O_2_ by PSII to H_2_O_2_ (Mehler reaction). Overall for each oxygen molecule consumed by Mehler reaction four electrons were transported from water to PSI via the linear electron transport chain (2 H_2_O + 2 O_2_ ⇔ H_2_O_2_ + O_2_). To avoid cleavage of H_2_O_2_ by catalase that would disturb the strict electron to oxygen stoichiometry, 1 mM sodium azide was added to the reaction cocktail.

The activity of Cytb_6_f complex was determined using 25 mM reduced duroquinol (DQH_2_) as the electron donor (replaces plastoquinone) and 100 μM MV as the PSI electron acceptor. This is a measure of linear electron flow at its maximum capacity.

### Difference spectroscopy

The contents of PSII and Cytb_6_f complex were measured in isolated thylakoids incubated in 0.003% (w/v) β-dodecyl-maltoside by difference spectroscopic quantifications of Cytb_559_ and Cytb_6_, respectively (Kirchhoff *et al*., 2002). The spectra were recorded with a Hitachi U3900 spectrometer (spectral range, 540–575 nm; 2-nm slit width) and analysed as described (Kirchhoff *et al*., 2002). Difference absorption spectroscopy of photooxidizable Cytf and P_700_ in leaves was measured with home-built flash spectrometer (Kirchhoff *et al*., 2011). For measurements, the leaf material was incubated in 5 mM MV-soaked Kimwipes for 30 min in darkness. P_700_ redox signals were derived from absorption changes at 820 and 900 nm, while Cytf redox kinetics were derived from absorbance changes at 554, 545 and 572 nm (Kirchhoff *et al*., 2004).

### Measurement of pmf and _Δ_pH

Pmf in leaves was deduced from ECS_total_ signals at 520 nm corrected by 550 nm (Klughammer *et al*., 2013), measured with a home-built flash spectrometer (Kirchhoff *et al*., 2011). Pmf was determined at the end of a 10 min illumination period (500 μmol photons m^−2^ s^−1^). Light-induced ΔpH was measured in freshly isolated thylakoids by 9-AA fluorescence quenching (Schuldinger *et al*., 1972). The magnitude of the ΔpH across thylakoid membranes was calculated from 9-AA quenching as described (Van *et al*., 1987). Measurements were performed with a HoribaYvon Fluoromax 4 spectrofluorometer (λ_ex_ = 400 nm; bandwidth: 2 nm; λ_em_ = 455 nm; bandwidth: 5 nm) with thylakoids (20 μg Chl ml^-1^) suspended in measuring buffer containing 6 μM 9-AA and 50 μM MV under continuous stirring. ΔpH was induced by red light of 1,000 μmol m^−2^ s^−1^. 1 μM of nigericin was added after each light cycle to obtain maximum fluorescence value during darkness.

### Protein analysis

Electrophoretic separation of proteins was carried out in 12 % (w/v) acrylamide SDS-gels. Following electrotransfer to PVDF membranes, proteins were identified by western blotting. Following antibodies from Agrisera (Vännäs, Sweden) were used: TIP2;1, TIP1;1/TIP1;2, actin, V-ATPase, H^+^-ATPase, Tic40, D1, Lhcb2 and RbcL. The anti-GFP antibody was obtained from Roche Diagnostics (Indianapolis, IN, USA).

### Accession numbers

Sequence data from this article can be found in the EMBL/Gen-Bank data libraries under accession numbers *At2g36830* (TIP1;1), *At3g26520* (TIP1;2) and *At3g16240* (TIP2;1).

## ACKNOWLEDGEMENTS

We are very grateful to the Centre for Cellular Imaging (Sahlgrenska Academy, Gothenburg University) for the use of confocal microscopes and helpful assistance. This work was supported by the Swedish Research Council and the Enkvists Foundation (C.S.). H.K. received support from the National Science Foundation (NSF-MCB115871), the United States-Israel Binational Agricultural Research and Development Fund (BARD US-4334-10), the US Department of Agriculture (ARC grant WNP00775), and Washington State University. This work was also supported by the French Ministère de l’Enseignement Supérieur et de la Recherche and INRA (K.B., B.S.) and University of Le Mans (B.S.). Travel Grants were provided by Helge Ax:Son Johnsons Foundation (A.B.), Adlebertska stiftelsen and the University of Le Mans (Grant: Chercheur haut niveau) (C.S.).

## SUPPORTING INFORMATION

### Supplemental Methods

Construction of TIP-GFP fusions and screening of *tip* mutants

## Supplemental Tables and Figures

**Table S1**. Levels of photosynthetic complexes in Col and *tip* mutants.

**Table S2**. Primers for GFP fusions PCR cloning.

**Table S3**. AGI codes, gene names and primer sequences for quantitative RT-PCR.

**Figure S1**. Confocal microscopic images of leaves from Col wild type plants and lines expressing TIP-GFP fusion proteins.

**Figure S2**. Growth of Ws plants and the *tip1;1* mutant.

**Figure S3**. Osmoregulatory volume changes in Ws and *tip1;1* mutant.

**Figure S4**. Light and osmoticum induced changes in isolated thylakoids.

**Figure S5**. Photosynthetic activity in the *tip1;1* mutant and Ws plants.

**Figure S6**. Western blot analysis of marker proteins for photosynthetic complexes.

**Figure S7**. Effect on intersystem transport and PSI.

**Figure S8**. Dynamics of photosynthesis and photoprotection in fluctuating light.

**Figure S9**. The effect of (NH_4_)(NO_3_) on the dynamics of photosynthesis and photoprotection in fluctuating light.

