## Supplemental methods for "TIP family aquaporins play role in chloroplast osmoregulation and photosynthesis"

### Construction of *TIP-GFP* fusion and plant transformation

The *Arabidopsis* lines expressing *TIP1;1* fused to GFP (*TIP1;1-GFP*) under the control of the endogenous *TIP1;1* promoter (~3 kb) were described in (Beebo *et al.,* 2009). To obtain the fusion constructs *TIP1;2-GFP* and *TIP2;1-GFP* under the control of the corresponding endogenous promoter, the genomic sequences of *TIP1;2* and *TIP2;1* starting ~3kb upstream of the translational start codon and ending before the stop codon were amplified by PCR (for primers see Table S2), cloned in the Gateway entry vector pDONR221 (Invitrogen) and transferred in front of the GFP sequence in the modified pMDC83 binary destination vector (Curtis and Grossniklaus 2003), in which the double 35S promoter was deleted. The binary vectors were introduced into *Agrobacterium tumefaciens* strain (GV3101) by heat shock method. The recombinant *Agrobacterium* clones were used to transform *Arabidopsis* plants through the floral dip method (Clough and Bent 1998). *TIP-GFP* transgenic plants were selected on MS agar plates containing 50 μg $ml^{-1}$ kanamycin.

### Screening of the *tip* mutants

The *Arabidopsis tip1;1* mutant is a T-DNA insertion mutant in Ws-4 background and was previously described (Beebo *et al.,* 2009). The *tip1;2* and *tip2;1* lines in Col-0 background contain an insertional Suppressor-mutator (dSpm), belong to the SLAT collection (Tissier *et al.,* 1999), and were obtained from the NASC stock center (accessions N116827 and N118785). To obtain the double mutant line *tip1;2/tip2;1*, the F2 generation of a cross between the two single mutant lines was screened by PCR to identify plants with a recombinant chromosome bearing the two mutations using the specific primers listed in Table S2. Quantitative real-time PCR using *UBQ10* as reference gene confirmed the reduction in the level of *TIP* mRNA in the next generation. Western blotting using anti-TIP antibodies confirmed the lack of the corresponding protein in the homozygous lines.
