## Supplemental tables and figures for "TIP family aquaporins play role in chloroplast osmoregulation and photosynthesis"

**Table S1** Levels of photosynthetic complexes in Col and *tip* mutants.

| Genotype | PSII Chl/$Cytb_{559}$[a] | $Cytb_6$ Chl/$Cytb_6$[a] | Cytf $A_{554}$-$A_{545}$-$A_{572}$[b] | PSI $A_{810}$-$A_{900}$[b] |
|---|---|---|---|---|
| Col | 281 ± 24 | 176 ± 10 | 0.0117 ± 0.0003 | 0.0047 ± 0.0001 |
| *tip1;2* | 318 ± 15 | 194 ± 18 | 0.0109 ± 0.0004 | 0.0044 ± 0.0004 |
| *tip2;1* | 332 ± 30 | 206 ± 26 | 0.0117 ± 0.0004 | 0.0047 ± 0.0002 |
| *tip1;2/2;1* | 306 ± 2 | 218 ± 22 | 0.0115 ± 0.0006 | 0.0044 ± 0.0002 |

Plants were grown for 6 weeks and dark-adapted overnight before the measurements.
[a] PSII and Cytb6f contents were measured by difference spectroscopic quantification of $Cytb_{559}$ and $Cytb_6$, respectively.
[b] Cytf and PSI contents were measured in leaves treated with methyl viologen (1 mM) for 40 min in darkness. Cytf redox level was monitored following absorption changes at 554 nm corrected by those at 545 and 572 nm, while PSI level was derived from absorption changes at 820 nm subtracted by those at 900 nm. The values are means ± SD (n = 5-7).

**Table S2** Primers for GFP fusions PCR cloning.

| AGI Code | Gene name | Primer name | Primer sequence (5'-3') |
|---|---|---|---|
| *At3g26520* | *TIP1;2* | attB1-S1 | GGGGACAAGTTTGTACAAAAAAGCAGGCTTCTTC AGTCGCTGTGTCCA |
| | | attB2-S1 | GGGGACCACTTTGTACAAGAAAGCTGGGTACCAG TAATCGGTGGTAGGCAAT |
| *At3g16240* | *TIP2;1* | attB1-D1 | GGGGACAAGTTTGTACAAAAAAGCAGGCTTCGAG AAAGATGCAAAGCAAA |
| | | attB2-D1 | GGGGACCACTTTGTACAAGAAAGCTGGGTAGAAA TCAGCAGAAGCAAGAGGA |

Primers for TIP1;1 were according to (Beebo *et al.,* 2009).

**Table S3** AGI codes, gene names and primer sequences for quantitative RT-PCR.

| AGI Code | Gene name | Forward primer sequence | Reverse primer sequence | amplicon size (bp) |
|---|---|---|---|---|
| *At4g05320* | *UBQ10* | CACACTCCACTTGGTCTT GCGT | TGGTCTTTCCGGTGAGAGTCT TCA | 71 |
| *At3g26520* | *TIP1;2* | ATTCCAGCGTTCGGTCTC TCTG | GGCGATTGGTGCGATTGTT | 147 |
| *At3g16240* | *TIP2;1* | TCGGACGCTGCTCTTGAT ACAC | AAAGTGACGGCTGGGTTCACA T | 125 |

Primers for TIP1;1 were according to Beebo *et al.,* (2009).

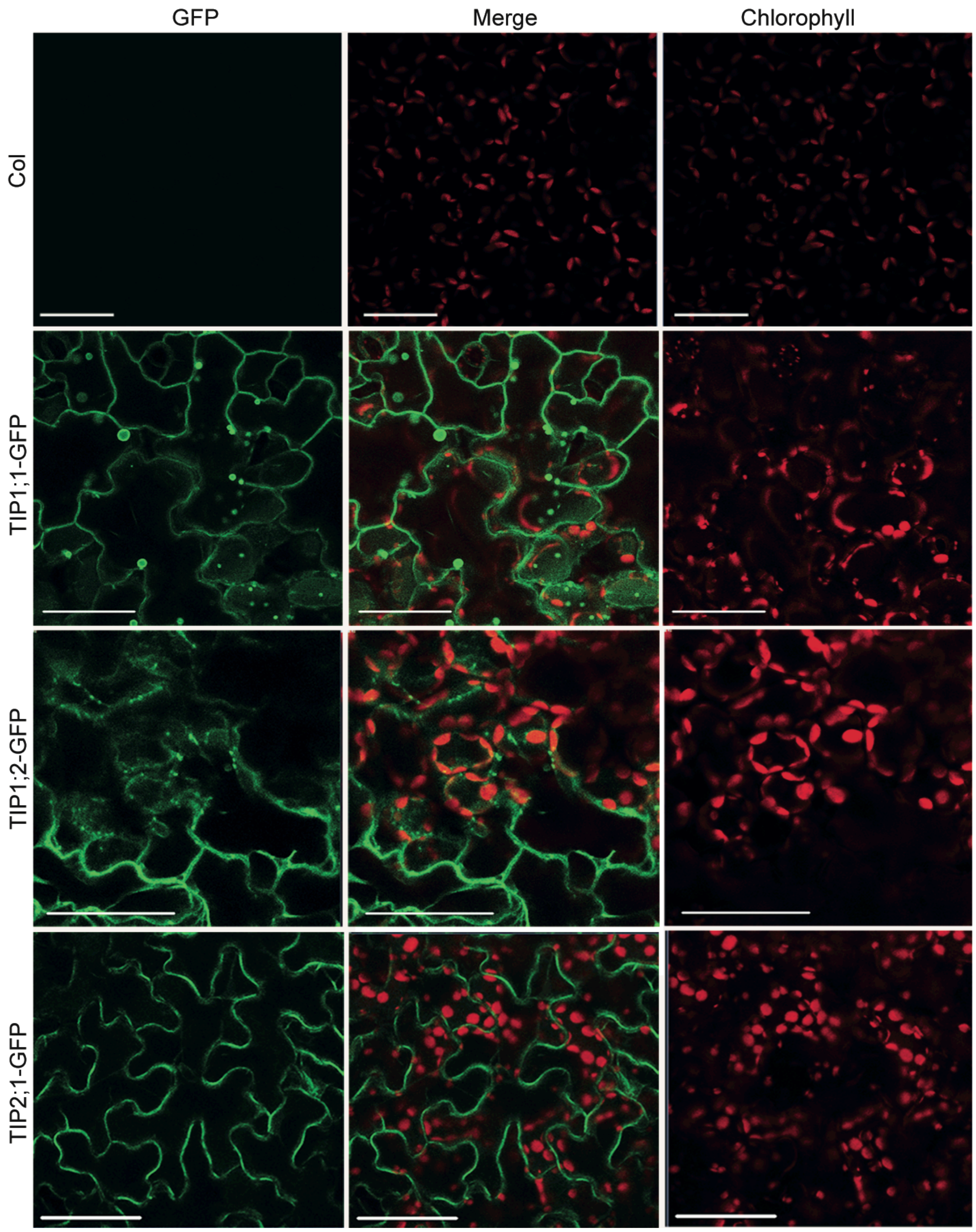


**Figure S1.** Confocal microscopic images of leaves from Col wild type plants and lines expressing TIP-GFP fusion proteins. The tonoplast was labeled by GFP in the leaves from all three transgenic lines. Chloroplasts were labeled by chlorophyll red autofluorescence but not by GFP at this magnification. Col leaves show only chlorophyll autofluorescence. Scale bar = 50 µm.

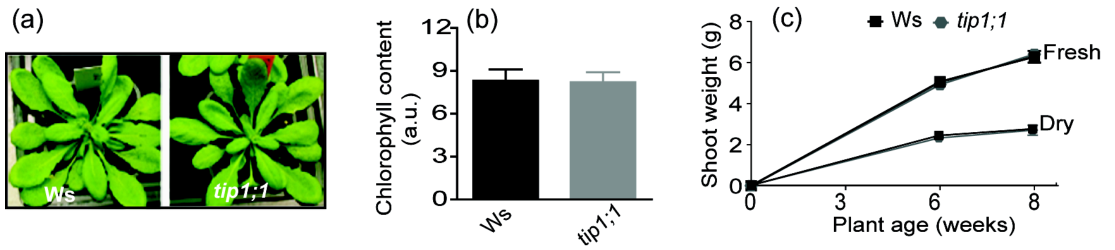


**Figure S2.** Growth of Ws plants and the *tip1;1* mutant.

(a) Representative photos at 8 weeks under 120 μmol photons $m^{-2}$ $s^{-1}$.

(b) Chlorophyll (Chl) content index at 8 weeks (n = 10-15).

(c) Shoot fresh and dry weight as a function of plant age.

The plotted values in all panels are means ± SD. No significant differences were observed between the mutant and Ws (Student's t-test, $P > 0.05$).

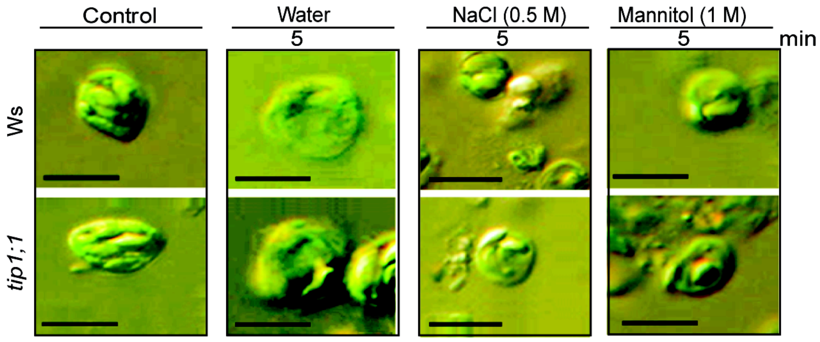


**Figure S3**. Osmoregulatory volume changes in Ws and the *tip1;1* mutant.

Light microscopic images of Percoll gradient chloroplasts incubated for 5 min in 0.3 M sucrose buffer (control), in water, 0.5 M NaCl or 1 M mannitol. Scale bar = 5 µm.

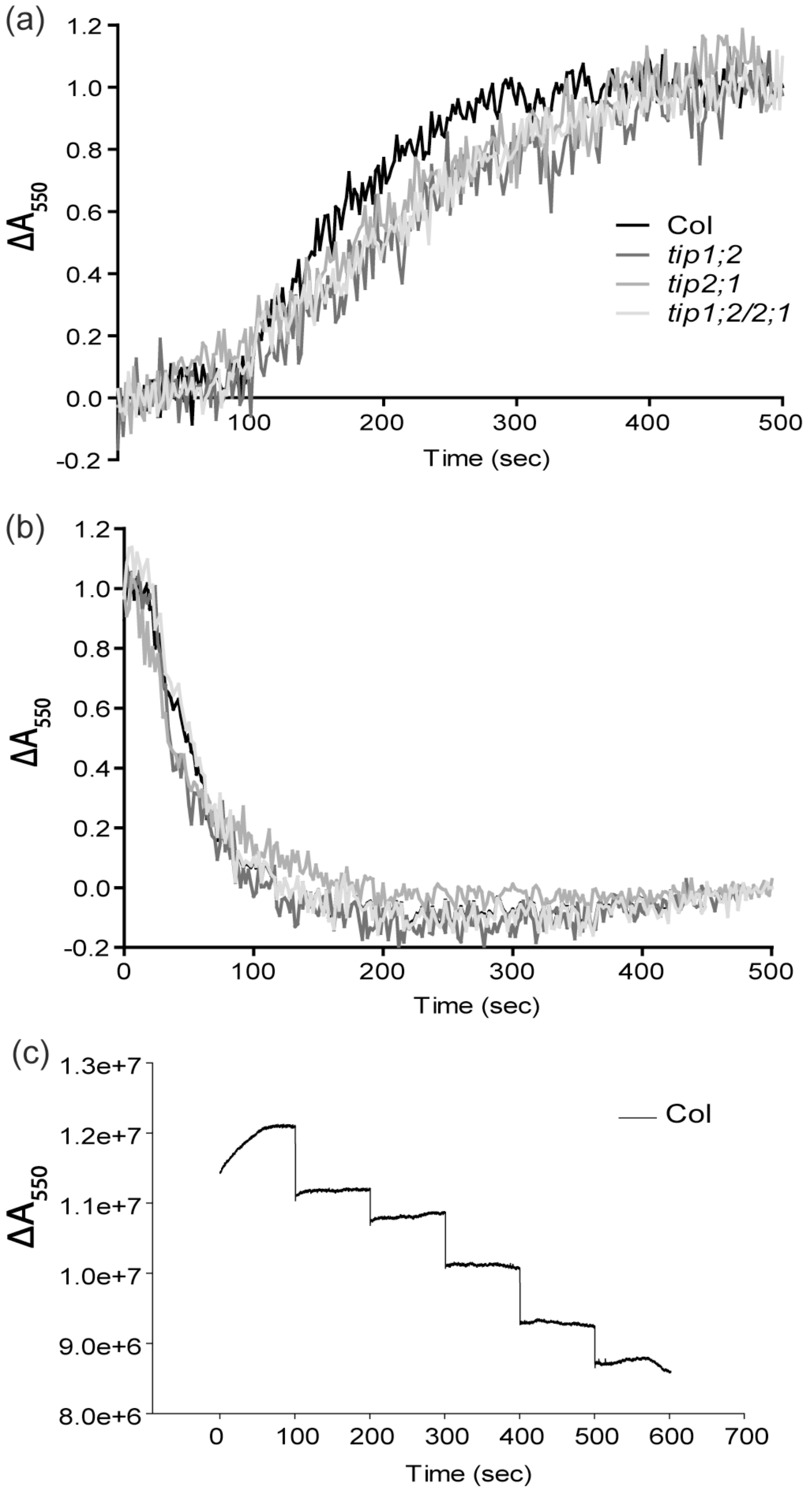


**Figure S4.** Light and osmoticum induced changes in isolated thylakoids.

(a) Light induced changes for Col and *tip* mutants.

(b) Relaxation in darkness following light-induced changes as in (a).

(c) Osmoticum induced scattering changes for Col thylakoids.

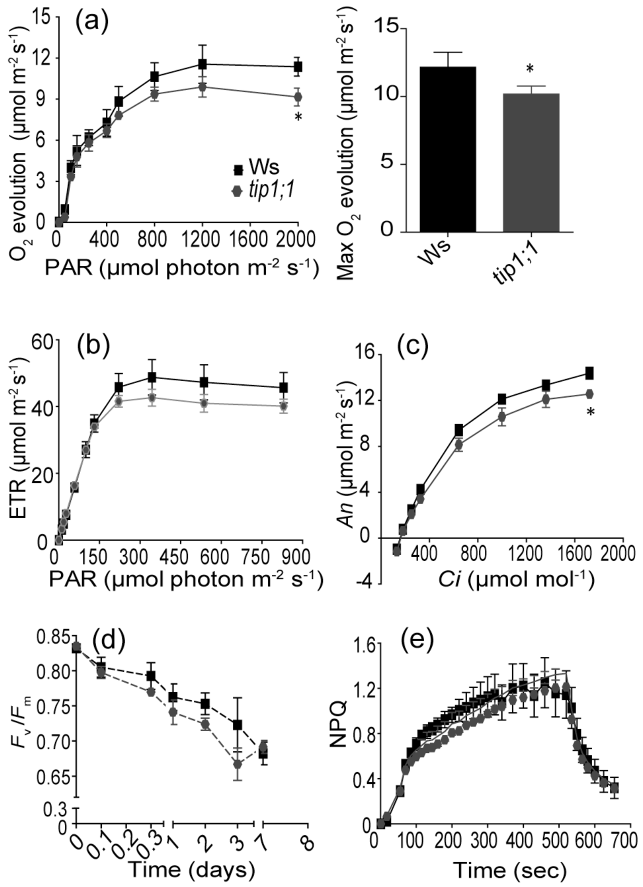


**Figure S5.** Photosynthetic activity in the *tip1;1* mutant and Ws plants.

(a) *Left:* Light response curves of $O_2$ evolution rates recorded in leaf discs (n = 3-5). *Right:* Maximal $O_2$ evolution from fitted light response curves.

(b) Light response curves of electron transfer rate (ETR) in intact plants (n = 10).

(c) Net $CO_2$ assimilation rate ($A_n$) versus intercellular $CO_2$ concentrations ($C_i$) recorded in leaf discs using light of 800 µmol photons $m^{-2}$ $s^{-1}$ (n = 3-4).

(d) $F_v/F_m$ of intact plants as a function of number of days in high light conditions (800 µmol photons $m^{-2}$ $s^{-1}$) (n = 5).

(e) Slow kinetics for induction of non-photochemical quenching (*NPQ*) formation recorded on intact plants using light of 800 µmol photon $m^{-2}$ $s^{-1}$ (n = 3-4). The plotted values are means ± SD (a-e). Significantly different values for parameters in the *tip1;1* mutant relative to Ws are indicated with asterisk (Student's t-test, $P < 0.05$).

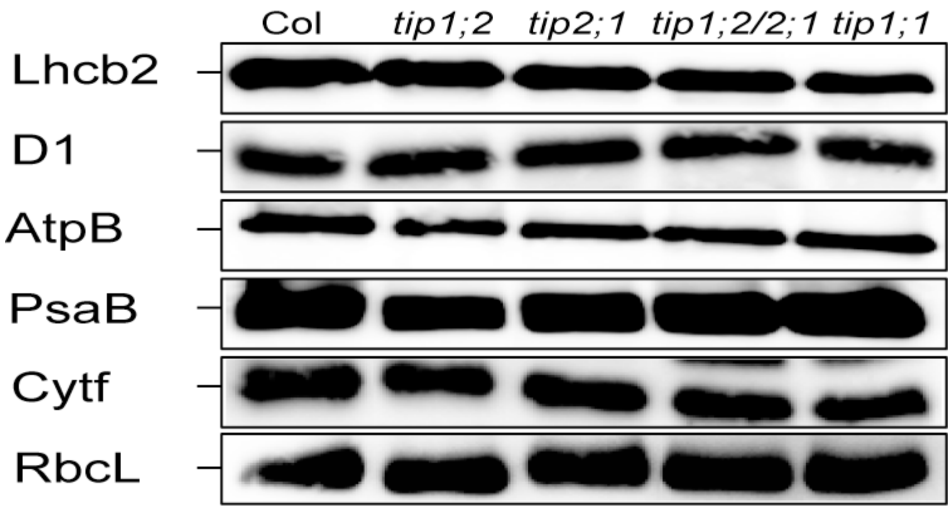


**Figure S6.** Western blot analysis of marker proteins for photosynthetic complexes. Lhcb2 and D1 (PSII), AtpB (ATP synthase). PsaA (PSI), and Cytf (cytochrome $b_6f$). RbcL is a subunit of stromal Rubisco.

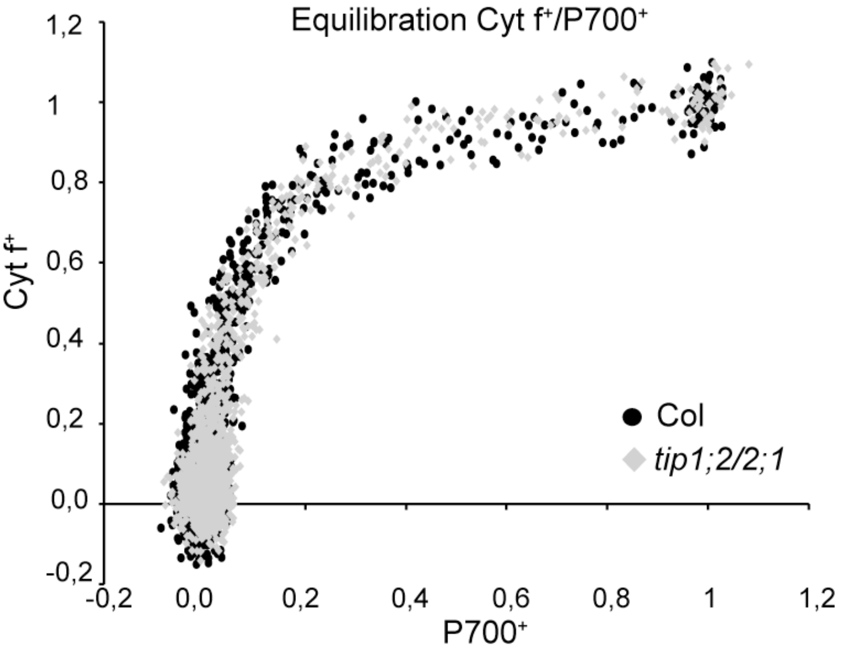


**Figure S7.** Effect on intersystem transport and PSI.

The redox kinetics of Cytf and $P_{700}$ were measured in leaves treated with methyl viologen (1 mM) for 40 min in darkness. Cytf redox kinetics were monitored following absorption changes at 554 nm corrected by those at 545 and 572 nm, while P700 kinetics were derived from absorption changes at 820 nm subtracted by those at 900 nm (n = 5).

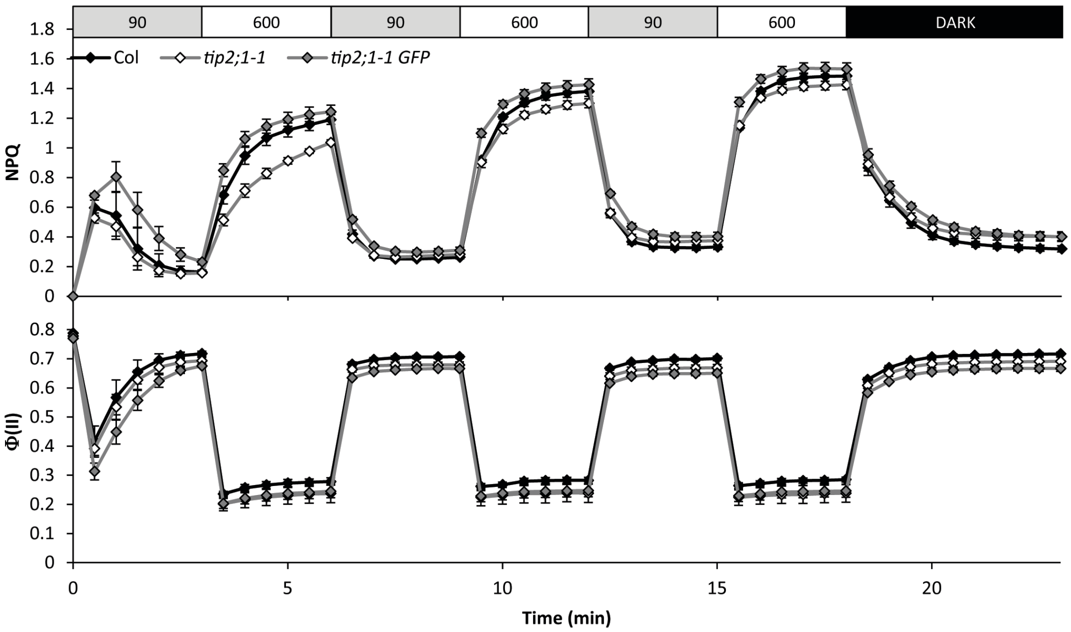


**Figure S8.** Dynamics of photosynthesis and photoprotection in fluctuating light.

Attached leaves 30-min dark adapted were exposed to fluctuating low light and high light. Slow kinetics for induction of non-photochemical quenching (*NPQ*) formation and photosystem II quantum yield Φ(II) were recorded. The plotted values are means ± SD (n=3-5).

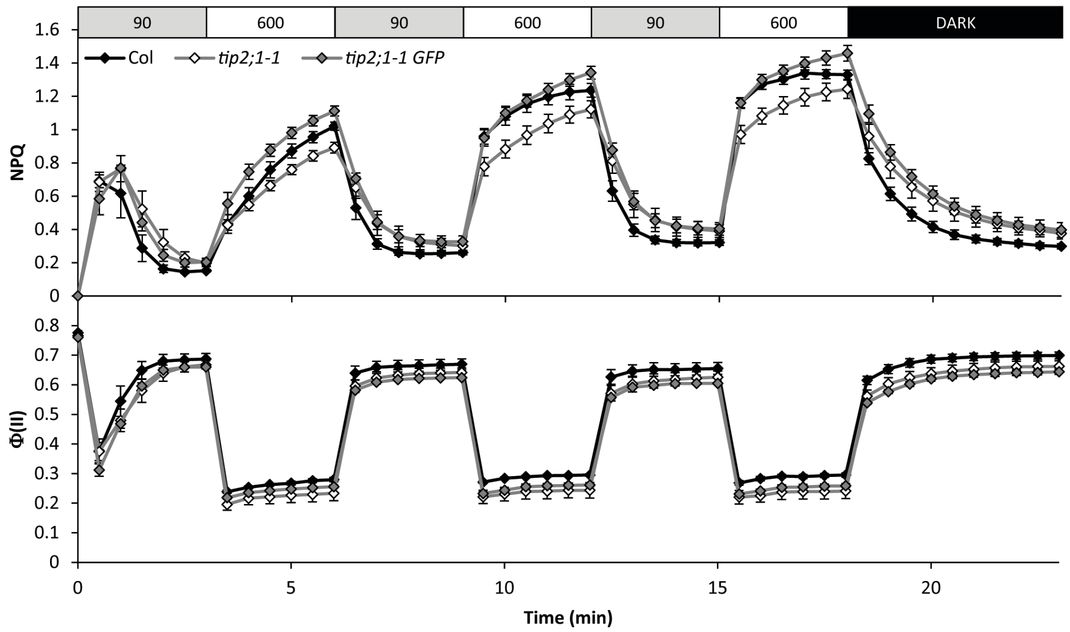


**Figure S9.** The effect of $(NH_4)(NO_3)$ on the dynamics of photosynthesis and photoprotection in fluctuating light.

Detached leaves were incubated for 30 min in growth light in 100 mM $(NH_4)(NO_3)$ followed by 30 min dark adaptation prior to recording the non-photochemical quenching (*NPQ*) and photosystem II quantum yield Φ(II) in fluctuating light. The plotted values are means ± SD (n=3-5).
